# Development of Metal organic Framework Based Biosensor to Detect the Coronavirus (Covid-19)

**DOI:** 10.1101/2022.08.20.504642

**Authors:** Hanoof Fahad Alkhodairy, Muhammad Naeem, Aasif Helal, Amjad Bajis Khalil

**Affiliations:** Department of Bioengineering, King Fahd University of Petroleum and Minerals (KFUPM), Dhahran 31261, Saudi Arabia; Interdisciplinary Research Centre for Hydrogen and Energy Storage, KFUPM, Dhahran 31261, Saudi Arabia

## Abstract

Recent outbreak of novel coronavirus (COVID-19) caused around 7 million deaths people worldwide and still afflicting on the global health, economy and social setup. Timely detection and diagnosis are crucial steps to reduce the spread and prevention of any pandemic. Different types of diagnosis methos has been used. In last decade nanomaterials and metal organic frameworks (MOFs) based biosensors has been developed to detect the other viruses. We have designed the Zeolitic imidazolate framework-8 (ZIF-8) based biosensor to detect the COVID-19. ZIF-8 work as fluorescence quenching and re-emergence platform to detect the COVID-19 RNA sequences. ZIF-8 platform is highly sensitive which can distinguish the highly conserved single strand RNA and with 200 pM concentrations. It can distinguish down to the single mismatch nucleotide in RNA sequences.

## 1. Introduction

SARS-CoV-2 virus cause infectious disease with severe respiratory symptoms called Coronavirus disease (COVID-19). The epidemic started from Wuhan, *China*, in December 2019. The WHO declared it pandemic on March 11, 2020. COVID-19 has drastically impacted the world’s demographics subsequent in more than 6 million deaths globally until June 2022.The coronavirus pandemic had a considerable impact on human life around the globe and presents an extraordinary challenge for the health care system, the food chain and the working hands. The social life limitations and the economic system disruption resulted by the pandemic is catastrophic; the world population is coping with the excessive poverty, the issue of undernourished disease shas been reached up to 690 million (WHO). [1] [2]

COVID-19 is currently being tested by two techniques: First, the Molecular tests, such as [polymerase](https://medlineplus.gov/lab-tests/pcr-tests/) [chain](https://medlineplus.gov/lab-tests/pcr-tests/) [reaction](https://medlineplus.gov/lab-tests/pcr-tests/) (PCR) tests that is a test to detect genetic material from a specific organism, such as a virus. A limitation to this test is that it requires an advanced laboratory and special equipment to be carried out. In addition to taking longer time in order for the polymerase to make many copies of the DNA. Ideally the samples are collected at a clinic, then sent to the laboratory. The second test is the Antigen tests (often referred to as rapid tests) in which the nucleocapsid protein (N protein) of SARS-CoV-2 from upper respiratory samples detected through immune essays. One disadvantage of antigen tests is that the tests have lower accuracy for people infected without symptoms. If a person gets a positive result ideally an RT-PCR test is performed afterward to confirm the positive result.

Recently, different types of nanomaterial based biosensors has been developed to detect DNA and RNA sequences. One of these methods include the use of Metal Organic Framework as a fluorescence quenching and emergence approach to identify the presence of nucleic acids (DNA/RNA) based on fluorophore-labelled oligonucleotide, fluorescence emitting organic molecule. [3] [4] A emerging unique type of nonporous materials, MOFs are substances are composed of metal ions and organic linker, MOFs have been drawing attention because of their ability of gasses separation and storage, catalysis, and biomedical applications [5, 6]

Zeolitic imidazolate frameworks (ZIFs) are a branch of the Metal Organic Frameworks that also have adjustable pore sizes and functional group functionality [7, 8]. In addition, Zeolitic imidazolate frameworks is stable in water due to its diverse structure nature. In this research work, (**Figure1**) we synthesized ZIF-8 based biosensor to generate a fluorescence assay for the recognition of Coronavirus conserved RNA sequences on DNA/RNA complementary base pairs basis. The general method used in this approach depended on the immersion of the fluorophore-labeled single-stranded probe DNA (P-DNA) to the Metal Organic Framework (Zif-8) forming an association with the ability of fluorescence quenching. Afterward, with the exact binding to the selected coronavirus RNA sequence, the formed double-stranded DNA/RNA becomes stronger and inflexible thus detach from the surface of the MOF resulting in the fluorescence re-generation. [1, 9]

**Figure 1:**
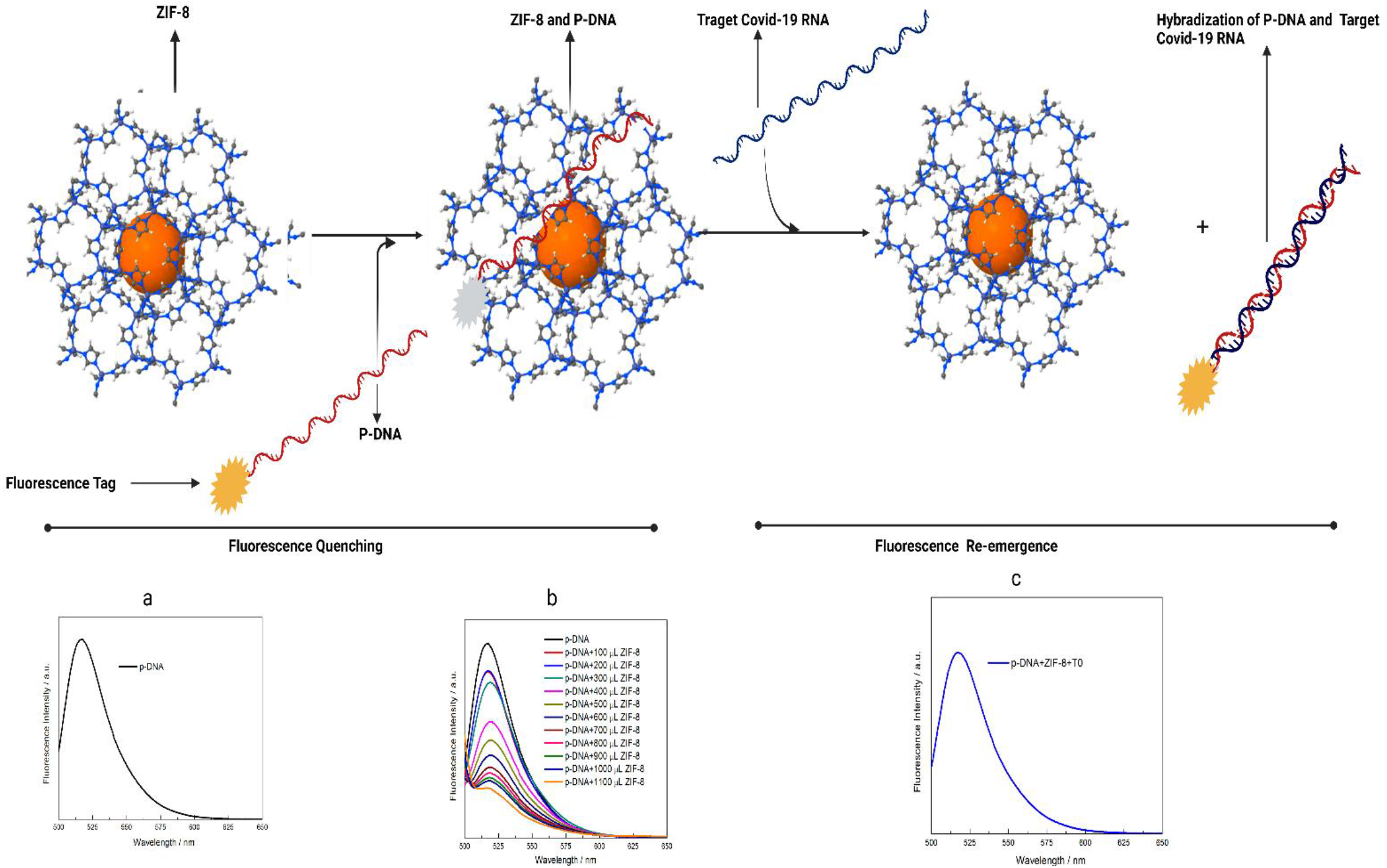
Schematic Representation of ZIF-8/P-DNA system as biosensor to detect the coronavirus (Covid-19) a) Fluorescence of Probe DNA (P-DNA). B) Fluorescence quenching of P-DNA by adding ZIF-8 C) Fluorescence re-emergence by adding the T0 Covid RNA into ZIF-8/P-DNA complex. (This figure has been created with Biorender.com)

## 2. Experiment Section

### 2.1. Materials

Zn(NO3)2•6H2O (98%, Aldrich), Co(NO3)2•6H2O (98%, Aldrich), and 2-methylimidazole (97%, Aldrich), were obtained from commercial sources and used without further purification.

The coronavirus (Covid-19) conserved sequences, 19 base pairs (bp) were determined through and Clustered Omega bioinformatic software further confirmation was done through NCBI BLAST against genome al possible strains of Covid-19 (Figure1) the sequence was unique in all strains of Covid-19.

All COVID-19 conserved sequences, Probe-DNA (P-DNA) and RNA with one and two mismatches were commercially from Microgen,inc, south korean biotechnology company. The sequence of Probe-DNA was (P-DNA) 5’-AGATGTCTTGTGCTGCCGGTA-3’ with 6-carboxyfluorescein (FAM)-labelled at 5′-terminal. The sequence of covid-9 RNA (T0) was, T0: 5’-UACCGGCAGCACAAGACAUCU-3’ is 100 percent complementary to the P-DNA sequences, With one mismatch T1: 5’-UACCGGCAGCAC**C**AGACAUCU-3’ and with two mismatches, T2: 5’-UACCGGCAGCAC**C**AGAC**G**UCU-3’. All the RNA sequences were suspended in Tris buffer solution placed at -80 C. The Probe-DNA was placed at 4 C. All the instruments employed for the development of of Covid-19 were autoclaved and disinfected.

The powder X-ray diffraction (PXRD) was carried out the Rigaku 2500 VBZ+/PC instrument. Thermogravimetric analysis (TGA) was carried out on a PerkinElmer Pyris Diamond TG Thermogravimetric Analyzer at a heating rate of 10 °C min−1.. [10]

### 2.2. Synthesis ZIF-8 MOF

#### Protocol

1. Weight 0.733 g Zn(NO3)2•6H2O (2.46 mmol)
2. Dissolved in 50 mL de-ionized water.
3. Weight 1.622 g HMe-Im (19.75 mmol)
4. Weight 2.00 g TEA (19.76 mmol) Add to 50 mL DI water and stirred up to cleared solution.
5. Add the Zn solution to the stirred HMe-Im/TEA solution.
6. the solution instantly became opaque white.
7. Further this was stirred for 10 minutes.
8. Centrifugation was done to remove the supernatant; the other remaining solid portion was mixed again with de-ionized H2O.
9. We placed it overnight and next day the centrifugation done one time with DI water and second time with ethanol. The tubes were dehydrated in oven at 110 °C and powder was selected.
10. Finally, the prepared solid was dried under vacuum at 150 °C for 1 to 1 and half hour.

### 2.3. TE Buffer

(TE buffer is very efficient buffer which protect the degradation of nucleic acids (RNA/DNA)in protected solution)

#### Protocol

1. Autoclave the flask.
2. Weight: 1.576g of Tris. 0.292g of EDTA
3. Add 80 ml Distilled water to the flask.
4. Add Tris and EDTA.
5. Bring to 100 ml with Distilled water.
6. Maintain Ph through HCL, and ph meter.
7. Autoclave and store at room temperature.

**Figure 2:**
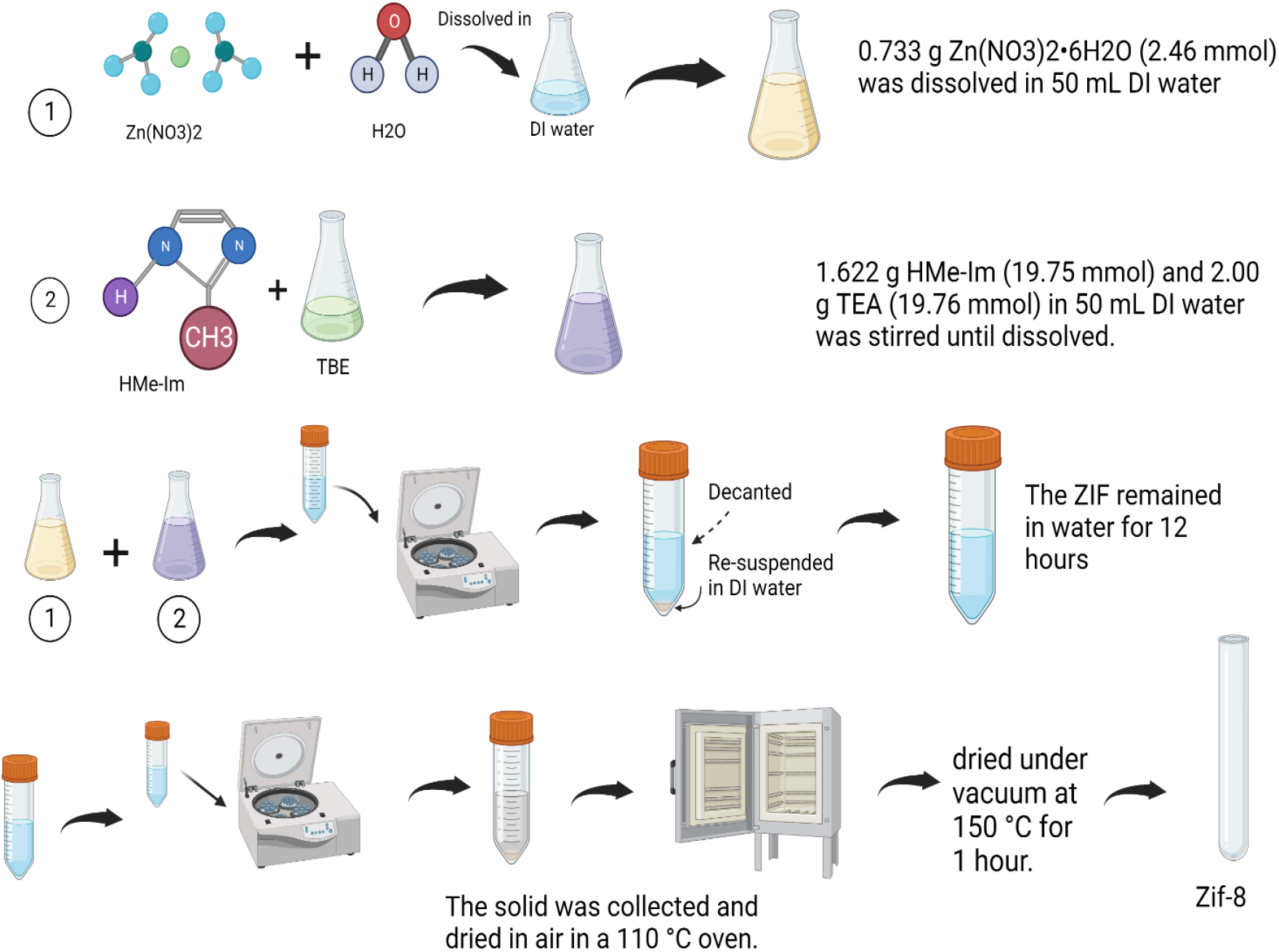
Schematic representation of ZIF-8 synthesis

### 2.5 Measurements

As the 1100 ul Zeolitic imidazolate frameworks 8 quenched the fluorescence of FAM-tagged P-DNA. The 900 uL of (100 nM) of FAM-tagged probe DNA (p-DNA) and 1100 uL of ZIF-8 (1mg/mL) ware added sequentially into the 2 ml tube after putting it 60 minutes at room temperature. The fluorescence was detected. Later on I ml from previous tube and 200 ul of targeted T0 COVID-19 RNA (200 pM) sequentially added into new 2mL tube. Incubate it on ice for 30 minutes and after it fluorescence was detected.

## 3. Results and Discussion

### 3.1. Characterization of synthesized Zeolitic imidazolate frameworks-8

ZIF-8 was synthesized with 79% yield from the reaction of zinc nitrate and 2-methyleimdazole in the de-ionized H2O at room temperature. The prepared ZIF-8 was further characterised through X-ray diffraction, and TGA, FT-IR and N2 adsorption. The IR data tells that the bands at 3135 and 2929 cm−1 are attributed to the aromatic and the aliphatic C-H stretch of the imidazole. The peak at 1584 cm−1 can be assigned as the C=N stretch mode specifically, whereas the intense and convoluted bands at 1350-1500 cm−1 are associated with the entire ring stretching. The bands in the spectral region of 900–1350 cm−1 are for the in-plane bending of the ring while those below 800 cm−1 are assigned as out-of-plane bending.

The ZIF-8 was dissolved in TAE buffer with P-DNA and complementary T0 RNA sequences for 2 days. Later the -XRD was compared with simple ZIF-8 (**Figure 3**). There was no structural difference examined in the peaks of only ZIF-8 and ZIF-8+P-DNA+ T0. The TGA analysis of prepared ZIF-8 was done. The TGA analysis showed that the ZIF-8 is stable until 510 °C.

**Figure 3:**
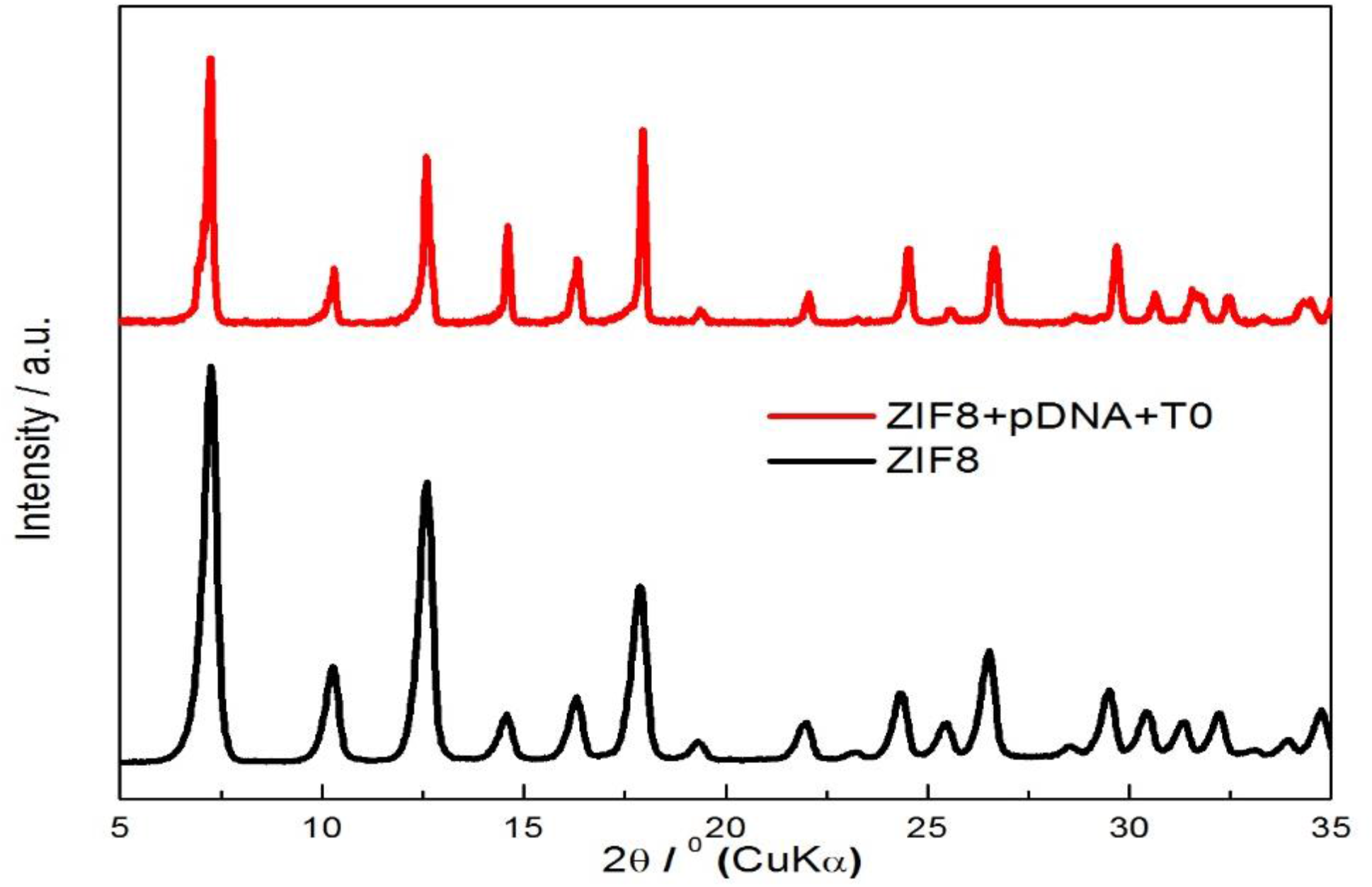
Black, The PXRD of ZIF-8. Red colour with PXRD of ZIF-+p-DNA+T0.

**Table 1:**
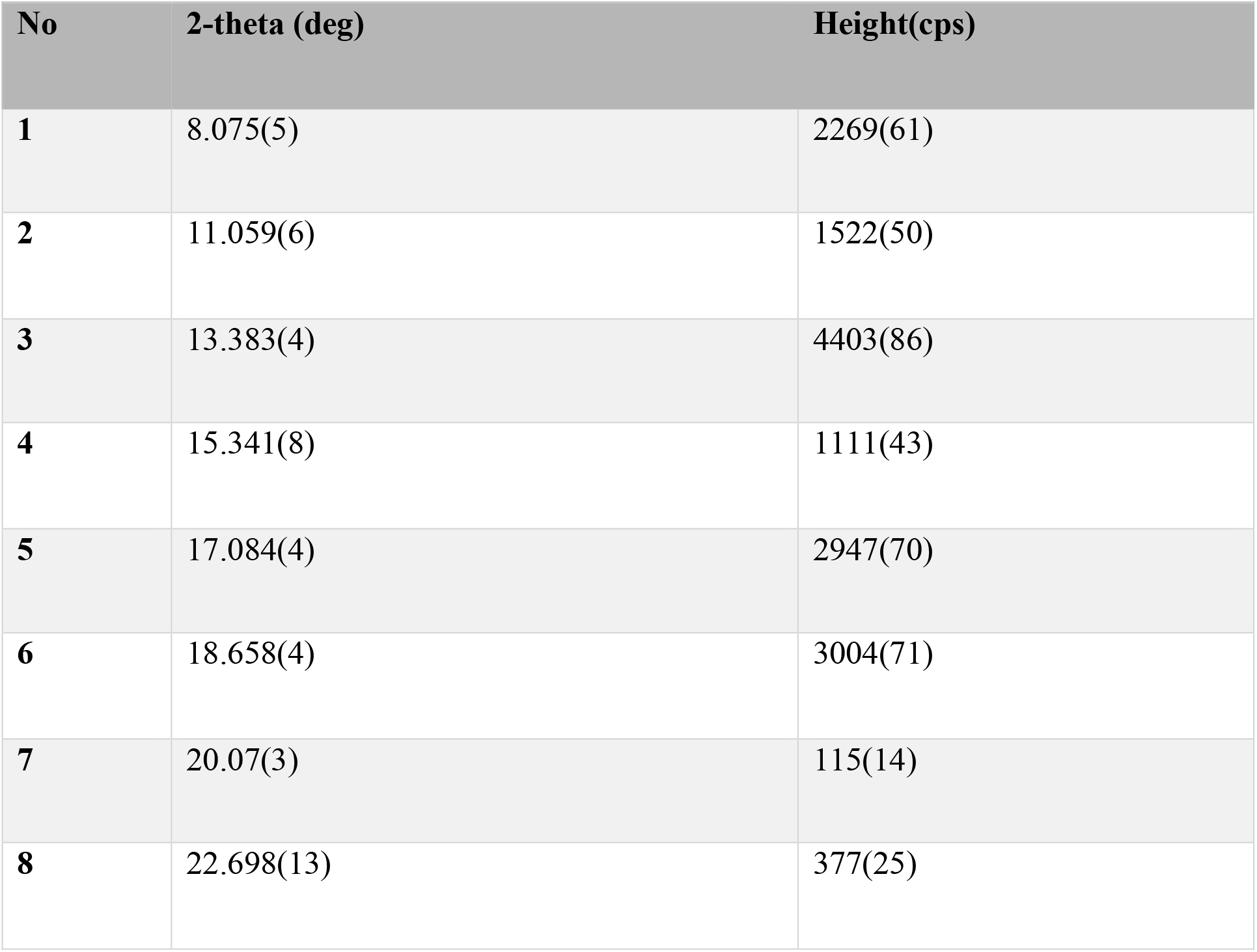
X-ray diffraction (XRD) analyses results

### 3.2. Biosensing Properties of ZIF-8 toward the Coronavirus (Covid-19) RNA sequences

As the ZIF-8 MOF has been synthesized with 2-methyleimdazole as a linker, Theoretically the imidazole contain the π-electron system which has ability to interact with the tagged DNA dye, fluorophore carboxyflurorescein (FAM) tagged at 5’ end of single strand P-DNA to form the π-π stacking that leads ultimately quenching the fluorescence. To test this we sequentially added the ZIF-8 to the solution of FAM containing PDNA (Figure 3), FAM 5’ AGATGTCTTGTGCTGCCGGTA-3’. It gradually decreases the fluorescence of P-DNA until the 1100 ul of 1mg/L ZIF-8 into the I mL of P-DNA solution in TAE buffer. Through experiments its showed that the complete quenching takes 60 minutes. This was proved the quenching of fluorescence was done through the π-π stacking.

To investigate the further quenching process, the P-DNA TAE buffer solution was tested individually against the zinc salt, zin nitrate and linker, 2-methyleimidalzole (Figure 4). The linker, 2-methyleimidazole effect on fluorescence quenching was negligible. But in contrast to linker the zinc nitrate quenched the fluorescence significantly higher than ZIF-8. This results indicate that the Zn(No3)2 can dimmish the fluorescence of P-DNA. Because of intercalation of zinc ion into the nucleotides of P-DNA or photoinduced electron transfer (PET) from the from fluorescence tag to Zinc ion. This also concluded that the ZIF-8, with 2-mthyleimdazole and Zinc ion can efficiently quench the fluorescence of P-DNA through π-π stacking and PET.

**Figure 4:**
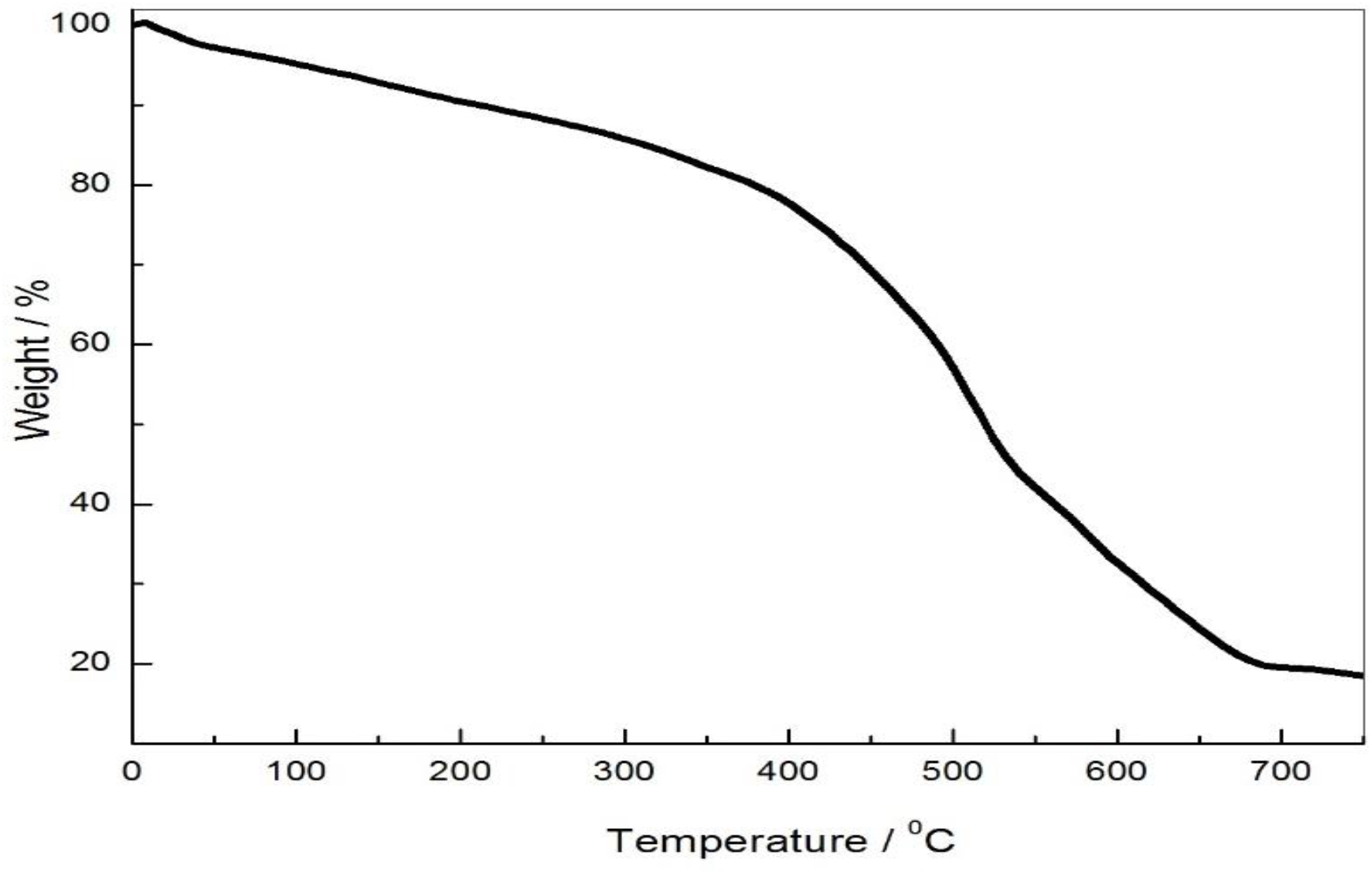
TGA analysis of prepared ZIF-8.

When the Coronavirus RNA, T0 sequence T0:5’UACCGGCAGCACAAGACAUCU-3’, complementary to P-DNA was added to the P-DNA/Zif-8 complex (Figure 5). There was 90 percent peak re-emergence. The re-emergence occurs due to the duplex formation of T0 RNA and P-DNA, RNA/DNA duplex that compulsion cause the P-DNA away from ZIF-8 [11]. But contrast to Zinc nitrate, there was a little re-emergence due to strong electrostatic interaction between the P-DNA and FAM. This shows that, the highly porous unique structure of ZIF-8/P-DNA complex is efficient that can quench the fluorescence and re-generate the fluorescence by adding T0 due to highest fluorescence quenching/ regeneration for T0 Covid-19 RNA sequence, the ZIF-8/P-DNA complex can be used a biosensor to detect the T0, conserved RNA sequences of Coronavirus.

**Figure 5:**
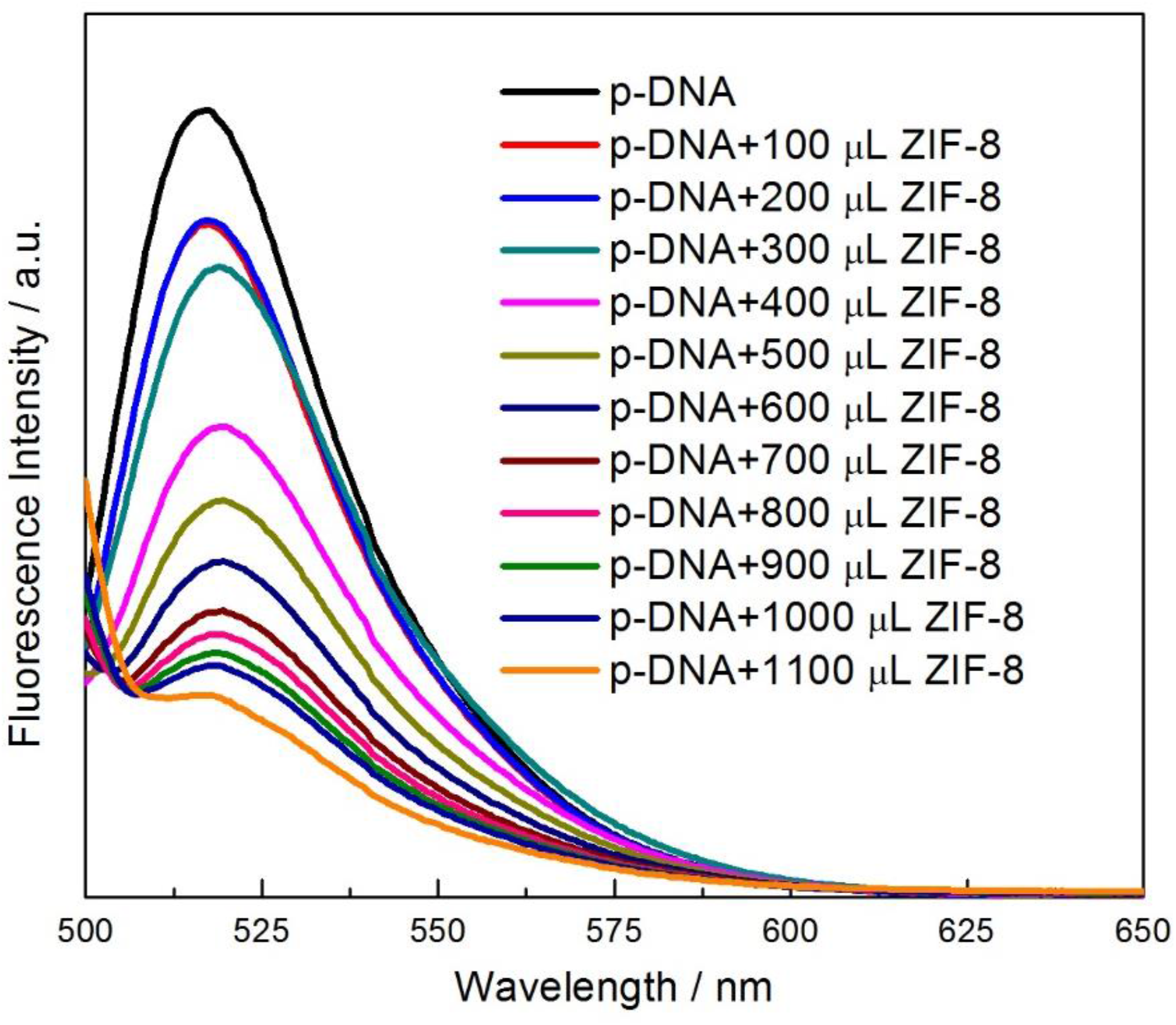
Gradually fluorescence quenching of P-DNA after adding the ZIF-8 sequentially.

**Figure 6:**
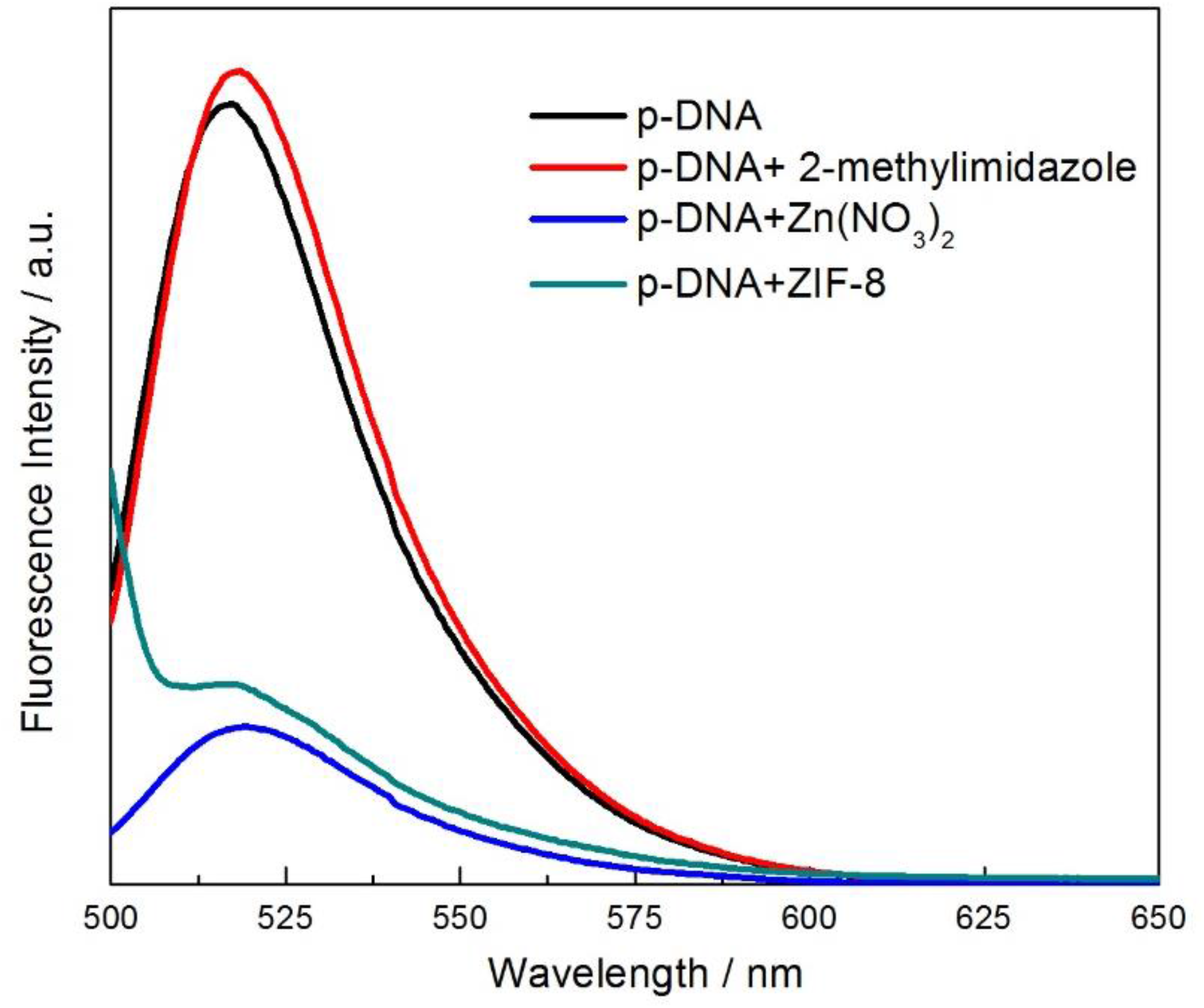
Fluorescence intensities of P-DNA (150 nM) by ZIF-8, 2-methyleimidazole, and zinc nitrate in 20 mM TAE buffer on at room temperature.

**Figure 7:**
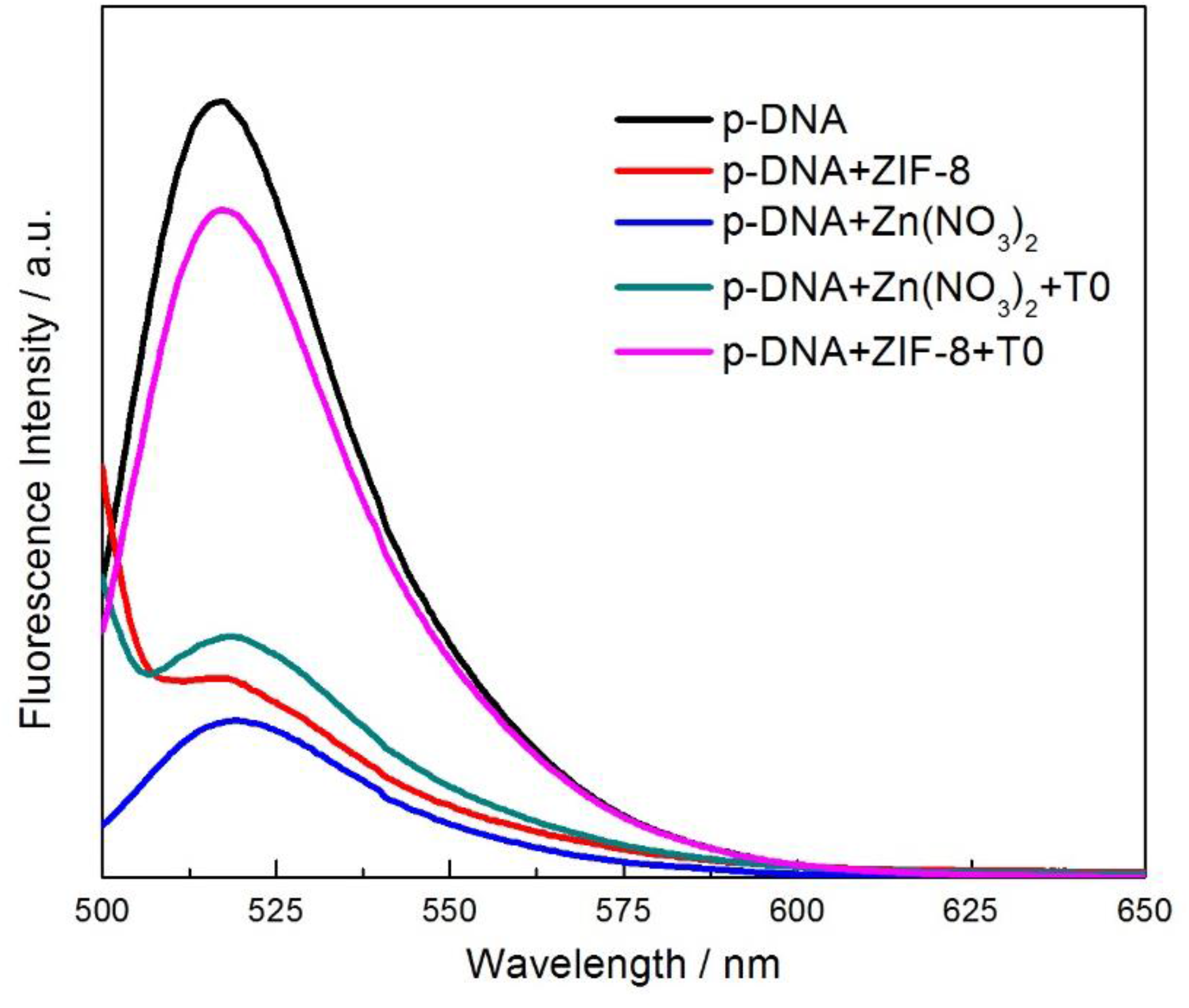
Fluorescence re-emergence of P-DNA/ Zif-8 with T0, P-DNA/ zinc nitrate with T0 in 20 mM TAE buffer on 4 °C at ice with different varying concentrations.

#### Single Mismatch Sensitivity

The further investigation was done of Zif-8/P-DNA complex biosensor toward the one mismatch T1, T1: 5’-UACCGGCAGCAC**C**AGACAUCU-3’ and with two mismatches, T2: 5’-UACCGGCAGCAC**C**AGAC**G**UCU-3’ to hybridize with the P-DNA **(Figure 8)**. The fluorescence emergence of T1 and T2 was 30 and 20 percent respectively to T0 with 90 percent. It shows that the ZIF-8/P-DNA complex could be used as a best biosensor to detect the Covid-19 through DNA/RNA hybridization.

**Figure 8:**
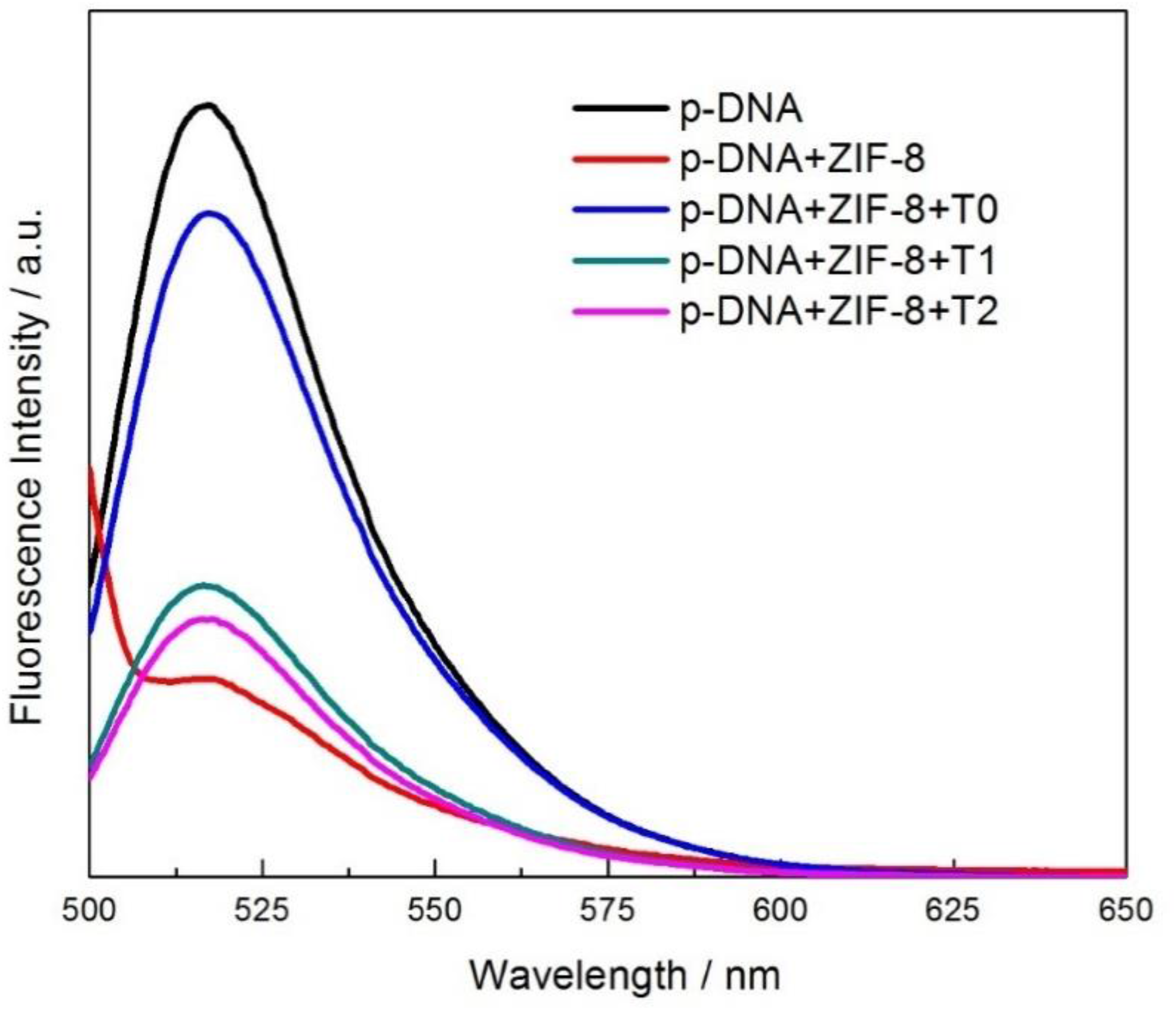
Fluorescence re-emergence of P-DNA/ Zif-8 with T0, P-DNA/ Zif-8 with T1, P-DNA/ Zif-8 with T2 with different varying concentrations.

#### Conclusion and Future Prospects

We have successfully synthesized the ZIF-8/P-DNA based biosensor complex system which can detect the coronavirus (Covid-19) conserved RNA sequences through DNA/RNA hybridization. These results encourage to further design and test MOF based biosensor detect the RNA as well as DNA viruses efficiently. This systematic platform can be employed to detect the future pandemics viruses.

## References

1. Damborska, D., et al., Nanomaterial-based biosensors for detection of prostate specific antigen. Microchimica acta, 2017. 184(9): p. 3049–3067.

2. Nadar, S.S. and V.K. Rathod, Immobilization of proline activated lipase within metal organic framework (MOF). International journal of biological macromolecules, 2020. 152: p. 1108–1112.

3. Wei, X., et al., Fluorescence biosensor for the H 5 N 1 antibody based on a metal– organic framework platform. Journal of Materials Chemistry B, 2013. 1(13): p. 1812–1817.

4. Zhu, X., et al., Metal–organic framework (MOF): a novel sensing platform for biomolecules. Chemical Communications, 2013. 49(13): p. 1276–1278.

5. Eddaoudi, M., et al., Systematic design of pore size and functionality in isoreticular MOFs and their application in methane storage. Science, 2002. 295(5554): p. 469–472.

6. Morris, R.E. and P.S. Wheatley, Gas storage in nanoporous materials. Angewandte Chemie International Edition, 2008. 47(27): p. 4966–4981.

7. Park, K.S., et al., Exceptional chemical and thermal stability of zeolitic imidazolate frameworks. Proceedings of the National Academy of Sciences, 2006. 103(27): p. 10186–10191.

8. Ulu, A., Metal–organic frameworks (MOFs): a novel support platform for ASNase immobilization. Journal of Materials Science, 2020. 55(14): p. 6130–6144.

9. Yang, S.-P., et al., Lanthanum-based metal–organic frameworks for specific detection of Sudan virus RNA conservative sequences down to single-base mismatch. Inorganic Chemistry, 2017. 56(24): p. 14880–14887.

10. Pan, Y., S. Zhan, and F. Xia, Zeolitic imidazolate framework-based biosensor for detection of HIV-1 DNA. Analytical Biochemistry, 2018. 546: p. 5–9.

11. Azam, T., et al., Tuning the hydrophobicity of MOF sponge for efficient oil/water separation. Chemical Physics Impact, 2020. 1: p. 100001.

